# Identical Tau Filaments in Subacute Sclerosing Panencephalitis and Chronic Traumatic Encephalopathy

**DOI:** 10.1101/2023.02.02.526771

**Authors:** Chao Qi, Masato Hasegawa, Masaki Takao, Motoko Sakai, Mayasuki Sasaki, Masashi Mizutani, Akio Akagi, Yasushi Iwasaki, Hiroaki Miyahara, Mari Yoshida, Sjors H.W. Scheres, Michel Goedert

## Abstract

Subacute sclerosing panencephalitis (SSPE) occurs in some individuals after measles infection, following a symptom-free period of several years. It resembles chronic traumatic encephalopathy (CTE), which happens after repetitive head impacts or exposure to blast waves, following a symptom-free period. As in CTE, when present, the neurofibrillary changes of SSPE are concentrated in superficial cortical layers. Here we used electron cryo-microscopy of tau filaments from two cases of SSPE to show that the tau folds of SSPE and CTE are identical. Two types of filaments were each made of two identical protofilaments with an extra density in the β-helix region. Like in CTE, the vast majority of tau filaments were Type I, with a minority of Type II filaments. These findings suggest that the CTE tau fold can be caused by different environmental insults, which may be linked by inflammatory changes.

## INTRODUCTION

Subacute sclerosing panencephalitis (SSPE) is a fatal disorder of the central nervous system that occurs following infection with measles virus and manifests itself after a symptom-free period of several years (1). It occurs in approximately 1 in 75,000 cases of measles (2). The neuropathology of SSPE is characterized by severe nerve cell loss, demyelination, perivascular lymphocytic infiltrations and viral intranuclear inclusion bodies. In around 20% of cases, abundant neurofibrillary tangles are present in cerebral cortex and other brain regions (3,4). It has been reported that tangles are made of paired helical filaments like those from Alzheimer’s disease brains (5,6) and that they stain for abnormally phosphorylated tau and ubiquitin (7).

Tangle-bearing cases of SSPE have mostly a long disease duration (8) and the tauopathy has been inferred to result from diffuse brain inflammation triggered by infection with measles virus and not from a direct effect of the virus (9). Neurofibrillary tangles of SSPE show staining profiles that are consistent with those of 3R+4R tauopathies, such as primary age-related tauopathy (PART), Alzheimer’s disease (AD) and chronic traumatic encephalopathy (CTE) (10). Unlike PART and AD, but like CTE, the neurofibrillary tangles of SSPE are present in superficial cortical layers (9,11).

We previously used electron cryo-microscopy (cryo-EM) to determine the atomic structures of tau filaments from a number of neurodegenerative conditions, which has resulted in a structure-based classification of tauopathies (12). We showed that the tau filament folds of 3R+4R tauopathies separate into two groups, the first of which is formed by PART, AD, and familial British and Danish dementias (FBD and FDD), and the second by CTE. The neurofibrillary pathology associated with some cases of Gerstmann-Sträussler-Scheinker disease (GSS) also belongs to the first group (13). We now report that the structures of tau filaments from two cases of SSPE are identical to those of CTE. This suggests that the CTE tau fold can form in response to different environmental insults, which may be linked by inflammatory changes.

## MATERIALS AND METHODS

### Clinical history and neuropathology

We determined the cryo-EM structures of tau filaments from the frontal cortex of two individuals with SSPE. Case 1 was a male who developed measles when 1.5 years old; at age 8, he developed a speech disturbance, as well as eating and walking difficulties. He was diagnosed with SSPE based on clinical presentation, a history of measles infection and characteristic electro-encephalogram and cerebrospinal fluid abnormalities. Despite intensive antiviral treatment, his condition worsened progressively and he died aged 42, after having been on mechanical ventilation for 11 years. The brain was severely atrophic with a weight of 439 g. Neuronal rarefaction was extensive in cerebrum, brainstem and cerebellum and there was a severe loss of myelinated nerve fibres. The clinicopathological characteristics of SSPE case 2 with a brain weight of 735 g have been described [case 3 in (9)].

### Extraction of tau filaments

Sarkosyl-insoluble material was extracted from the frontal cortex of cases 1 and 2 of SSPE, as described (14), with minor modifications. Briefly, tissues were homogenized with a Polytron in 40 vol (w/v) extraction buffer consisting of 10 mM Tris-HCl, pH 7.4, 0.8 M NaCl, 10% sucrose and 1 mM EGTA. Homogenates were brought to 2% sarkosyl and incubated for 30 min at 37° C. Following a 10 min centrifugation at 27,000 g, the supernatants were spun at 257,000 g for 30 min. Pellets were resuspended in 2 ml extraction buffer containing 1% sarkosyl and centrifuged at 166,000 g for 20 min. The resulting pellets were resuspended in 100 μl phosphate-buffered saline (PBS) and used for subsequent analyses.

### Immunolabelling and histology

Immunogold negative-stain electron microscopy and immunoblotting were carried out as described (15). For immunoelectron microscopy, the samples were applied onto collodion membrane-applied mesh, blocked with 0.3% gelatin, incubated with AT8 (1:100) for 1 h at 37° C, followed by a 1 h incubation at 37° C with 10 nm gold-labelled secondary antibody (1:50) and staining with 2% phosphotungstate. For immunoblotting, samples were run on 5-20% gradient gels (Fuji Film). Proteins were then transferred to a polyvinylidene difluoride membrane and incubated with the following primary antibodies overnight at room temperature: Tau N (1:1,000), AT8 (1:500), RD3 (1:500), anti-4R (1:1,000), Tau354-369 (1:1,000) and T46 (1:1,000). Following washing in PBS, the membranes were incubated with biotinylated anti-mouse or anti-rabbit secondary antibody (Vector, 1:500) for 1 h at room temperature, followed by a 30 min incubation with avidin-biotin complex and colour development using NiCl-enhanced diaminobenzidine as substrate. Histology and immunohistochemistry were carried out as described (16). Brain sections were 8 μm thick and were counterstained with haematoxylin. Primary antibodies were: RD3 (1:1,000); anti-4R (1:1,000); AT8 (1:300); anti-measles virus antibody (1:1,000); Iba-1 (1:2,000); CD3 (1:1,000).

### Electron cryo-microscopy: Sample preparation and data collection

Extracted tau filaments were centrifuged at 3,000 g for 1 min and applied to UltrAuFoil cryo-EM grids (17), which were glow-discharged with an Edwards (S150B) sputter coater at 30 mA for 1 min. Aliquots of 3 μl were applied to the glow-discharged grids, blotted with filter paper and plunge-frozen into liquid ethane using a Vitrobot Mark IV (Thermo Fisher Scientific) at 100% humidity and 4°C. Cryo-EM images were collected on a Titan Krios electron microscope (Thermo Fisher Scientific) operated at 300 kV and equipped with a Falcon-4 direct electron detector. Images were recorded during 6 s exposures in EER (electron event representation) format (18) with a total dose of 40 electrons per A^2^ and a pixel size of 0.824 Â.

### Electron cryo-microscopy: Image processing

Image processing was performed using RELION-4.0 (19,20), unless indicated otherwise. Raw movie frames were gain-corrected, aligned, dose-weighted and summed into a single micrograph. Contrast transfer function (CTF) parameters were estimated using CTFFIND-4.1 (21). Filaments were picked manually and segments extracted initially with a box size of 1024 pixels. 2D classification was used to remove suboptimal segments and to separate Type I from Type II filaments. Selected class averages for Type I and Type II filaments were then re-extracted with a box size of 400 (for SSPE case 1) or 300 (for SSPE case 2) pixels. Initial models were generated *de novo* from 2D class averages using *relion_helix_inimodel2d* (22). Helical twist and rise were optimised during 3D auto-refinement. Bayesian polishing and CTF refinement (23) were used to improve the resolution of reconstructions of Type I filaments. Final maps were sharpened using standard post-processing procedures in RELION and their resolutions were calculated based on the Fourier shell correlation (FSC) between two independently refined half-maps at 0.143 (24). Helical symmetry was imposed on the post-processed maps using the *relion_helix_toolbox* program (25).

### Model building and refinement

Atomic models of published CTE filament structures (26) (PDB:6NWP; PDB:6NWQ) were docked manually in the density using Coot (27). Model refinements were performed using *Servalcat* (28) and Refmac5 (29,30). Models were validated with MolProbity (31). Figures were prepared with ChimeraX (32) and Pymol (33). Further details of data acquisition and image processing are given in Supplementary Table 1.

## RESULTS

For cryo-EM, we extracted tau filaments from the frontal cortex of two cases of SSPE. Both individuals had measles as children, with the clinical picture of SSPE manifesting itself following a number of symptom-free years. They belong to the minority of cases of SSPE with neurofibrillary lesions.

In SSPE case 1, immunohistochemistry with anti-tau antibodies showed abundant neurofibrillary tangles that were stained by anti-tau antibodies specific for 3R tau, 4R tau and phospho-tau (AT8) (Figure 1). Neurons and glial cells with intranuclear inclusion bodies were observed using an antibody against measles virus. Microglial cells were activated, as shown by Iba-1 staining; the same was true of cytotoxic and T helper lymphocytes, as evidenced by CD3 staining (Figure 1). Similar abnormalities have been described in SSPE case 2 (9).

**Figure 1.**
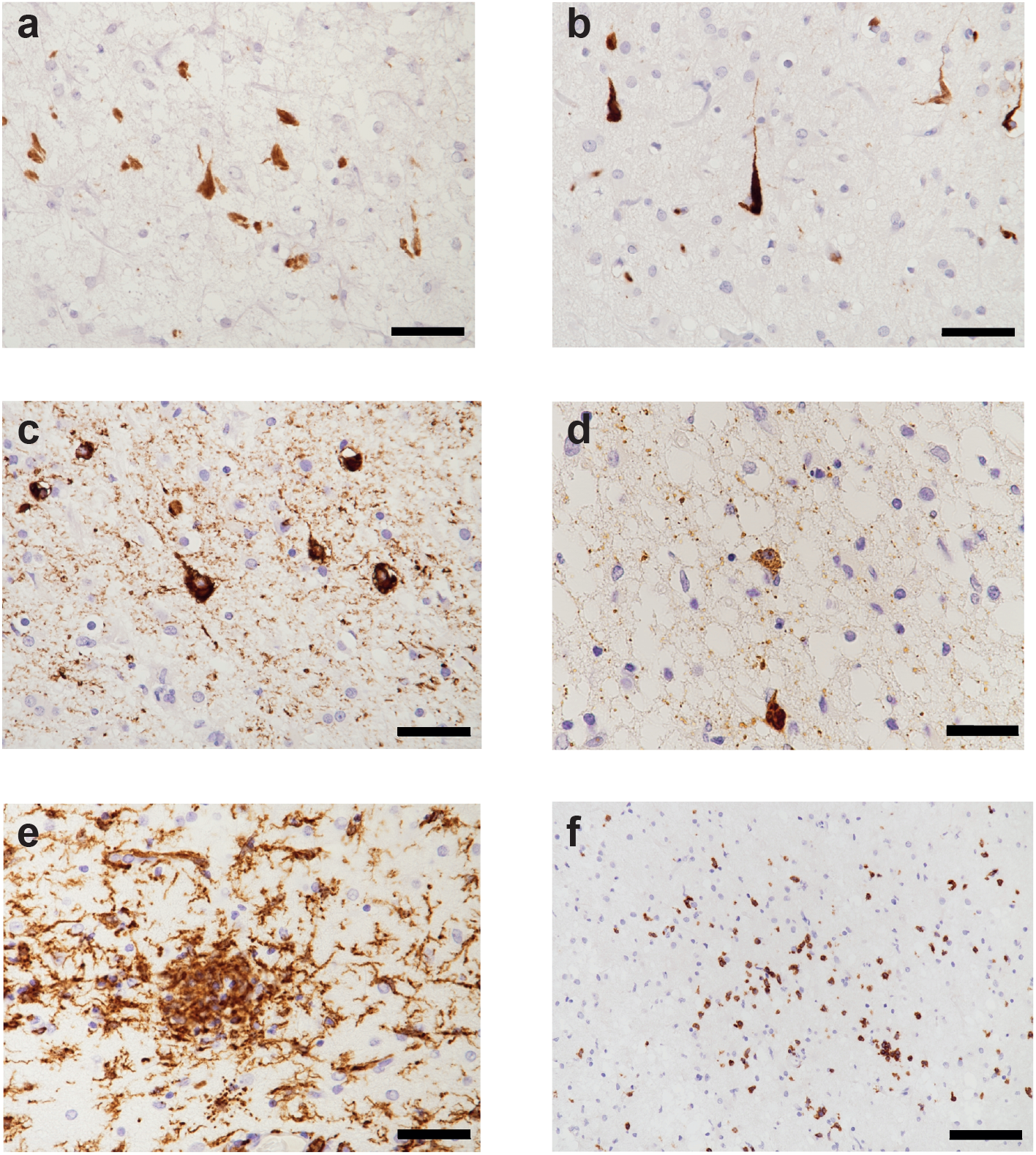
Frontal cortex from SSPE case 1: Immunohistochemical characterisation of tau inclusions and inflammatory changes. (a), RD3 (specific for 3R tau)-immunoreactive nerve cells and neuropil threads. (b), RD4 (specific for 4R tau)-immunoreactive nerve cells and neuropil threads. (c), AT8 (specific for pS202 and T205 tau)-immunoreactive nerve cells and neuropil threads. (d), Antibody against measles virus shows neuronal staining. (e), Iba-1-immunoreactive microglial cells. (f), CD3-immunoreactive lymphocytes. Scale bars: a-e, 50 μm; f, 100 μm.

Filaments from the sarkosyl-insoluble fractions were decorated by anti-tau antibodies and gave the same bands on Western blots as those from AD brains (Figure 2). It has previously been shown that the bands of sarkosyl-insoluble tau from AD are identical to those from CTE (34). The observed bands indicate that the filaments of SSPE cases 1 and 2 are made of all six tau isoforms in a hyperphosphorylated state, consistent with previous results (9).

**Figure 2.**
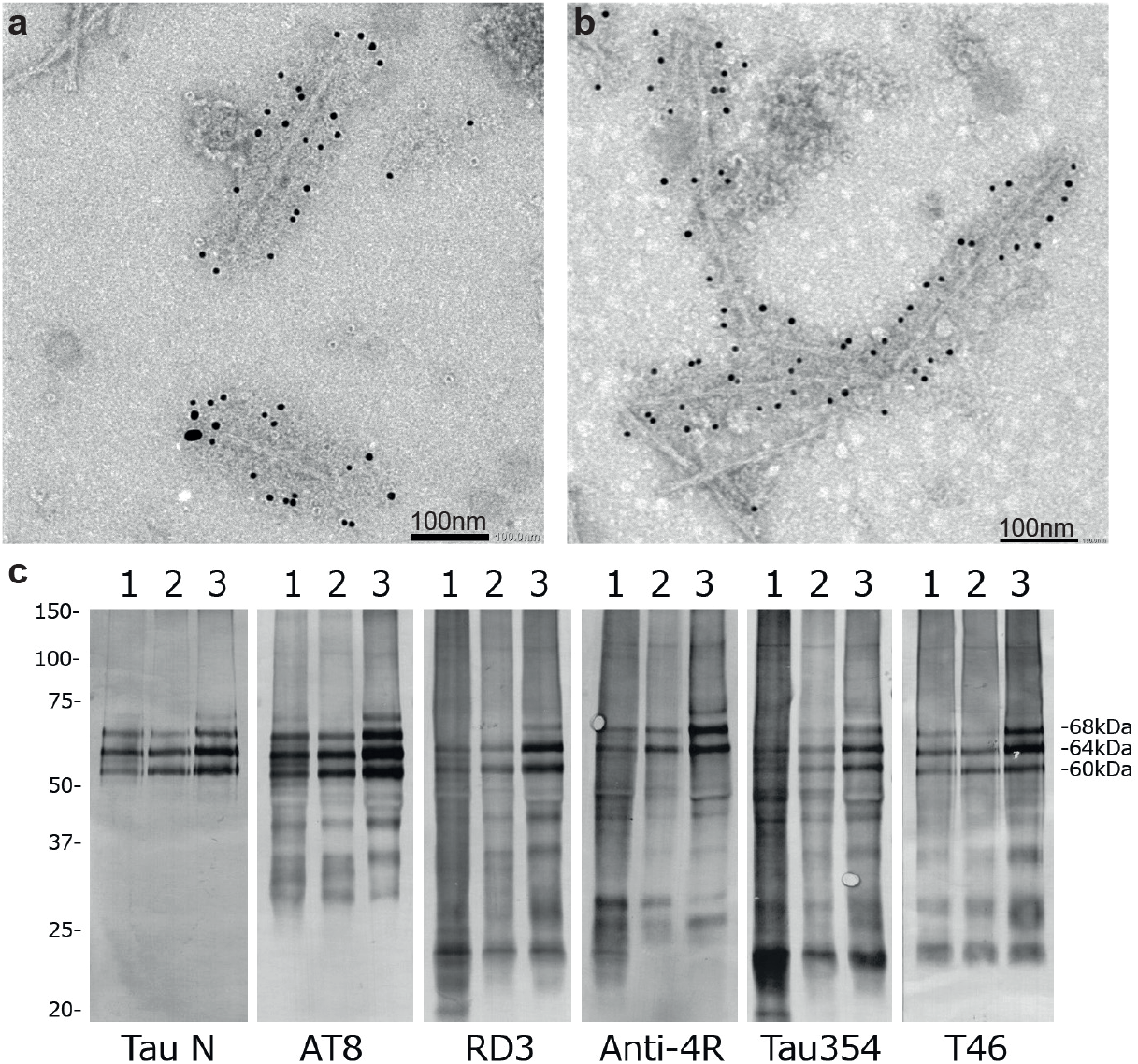
Immunolabelling and immunoblotting of tau filaments from SSPE. (a,b), Immunoelectron microscopy of filaments from SSPE cases 1 (a) and 2 (b) using anti-tau antibody AT8. (c), Immunoblotting of sarkosyl-insoluble fractions using anti-tau antibodies: Tau N; AT8; RD3; Anti-4R; Tau354; T46. Lanes: 1, SSPE case 1; 2, SSPE case 2; 3, AD.

By cryo-EM, we show that the CTE fold of assembled tau is characteristic of SSPE cases 1 and 2 (Figure 3; Supplementary Figure 1). As in the CTE fold, two types of tau filaments were present, each with an unknown, internal density in the β-helix of the structured core. Type I and Type II filaments are both made of two identical protofilaments, with different interprotofilament packing, i.e. type I and type II CTE filaments are ultrastructural polymorphs. Type I comprised more than 90% of the observed filaments and Type II less than 10%, similar to what has been observed in CTE (26).

**Figure 3.**
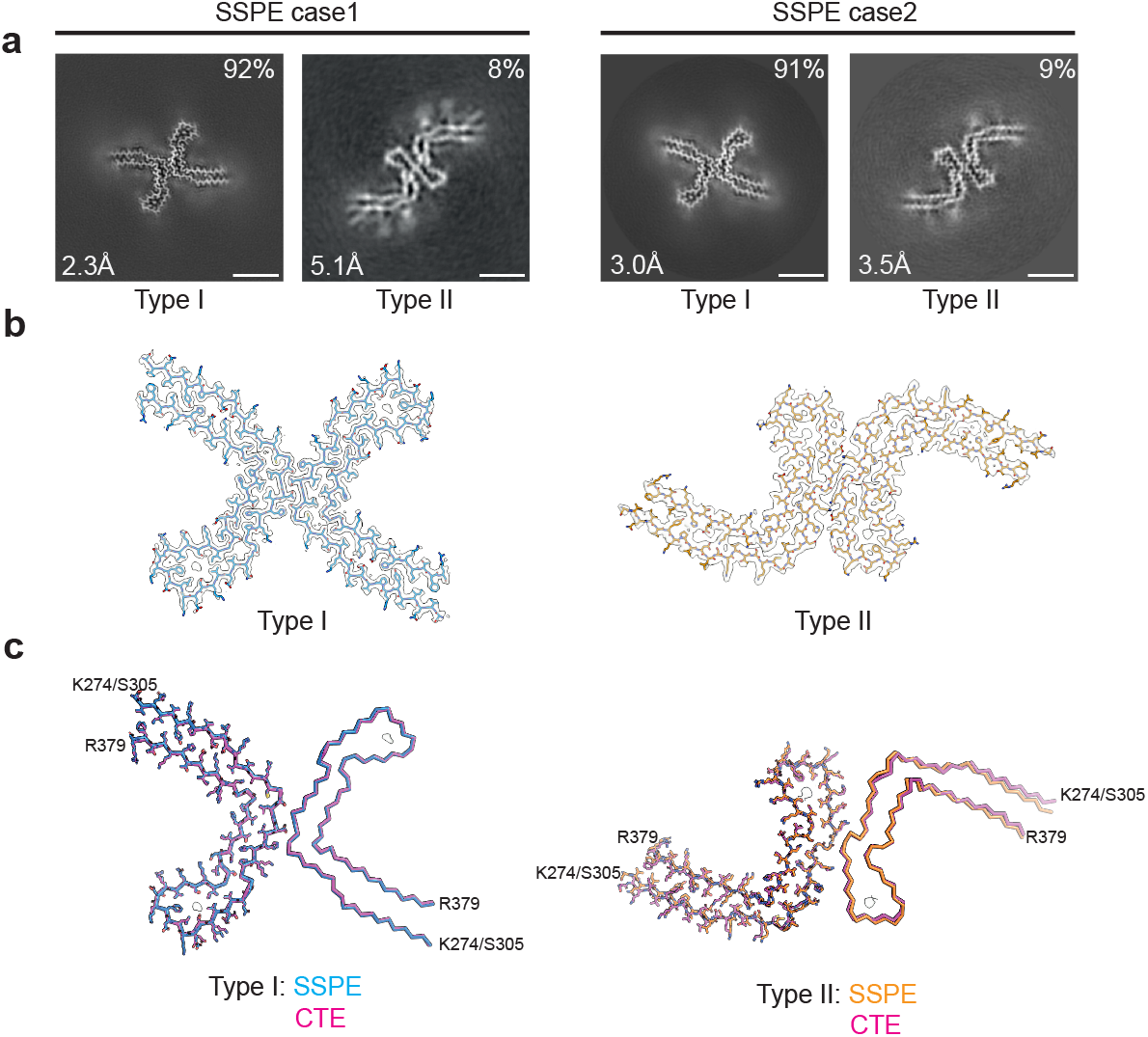
Cryo-EM cross-sections and structures of tau filaments from SSPE. (a), Cross-sections through the cryo-EM reconstructions, perpendicular to the helical thickness and with a projected thickness of approximately one rung, are shown for SSPE cases 1 and 2. A majority of Type I and a minority of Type II tau filaments are each made of two copies of a single protofilament arranged in different ways (ultrastructural polymorphs). Scale bar, 5 nm. (b), Cryo-EM density maps (grey transparent) of SSPE Type I and Type II tau filaments and the atomic models coloured blue (Type I) and orange (Type II). (c), SSPE Type I (blue) and Type II (orange) filaments overlaid with CTE Type I (magenta) and CTE Type II (magenta) filaments. The filament core extends from tau residues K274/S305-R379.

## DISCUSSION

SSPE and CTE share a long interval between the primary insult (measles virus infection and repetitive head impacts or blast waves) and the appearance of clinical symptoms. Like AD and CTE, tangle-bearing cases of SSPE are characterised by the presence of abundant filamentous inclusions made of all six brain tau isoforms. Unlike AD, CTE and tangle-bearing cases of SSPE share the formation of abundant tau inclusions in cortical layers II and III (9,11). We previously showed that the CTE fold differs from the Alzheimer tau fold by adopting a more open conformation of the β-helix region, which contains an internal density of unknown identity (26). In the presence of NaCl, recombinant tau comprising amino acids 297-391 assembles into filaments with the CTE fold, but in its absence, the Alzheimer tau fold forms (35).

We now find that the tau fold of SSPE is identical to that of CTE. As in CTE, two types of filaments, each made of two identical protofilaments, were present in SSPE cases 1 and 2. Western blots of sarkosyl-insoluble tau fractions indicate that these filaments are made of all six (3R+4R) brain tau isoforms (12,13,26,36). For 3R+4R tauopathies, one protofilament fold (the Alzheimer fold) is found in PART, AD, GSS, FBD and FDD, and the second protofilament fold (the CTE fold) is found in CTE. The present findings show that SSPE is a second example of a 3R+4R tauopathy with the CTE fold. It remains to be seen if other conditions with tau inclusions in cortical layers II and III also share the CTE fold.

Inflammation may be what CTE and SSPE have in common. In SSPE case 2, extensive inflammatory changes have been described, with perivascular lymphocyte infiltration, aggregates of hypertrophic astrocytes and activated microglia (9). Here we show that the frontal cortex from SSPE case 1 also exhibited microglial activation and lymphocyte infiltration. Immunoreactivity for measles virus was present in both cases of SSPE, even though they had undergone antiviral treatment. In untreated cases of long disease duration, measles virus was detected in cases with neurofibrillary lesions (37). In treated cases, the detection frequency of measles virus was decreased, even though neurofibrillary pathology was unaffected, suggesting that antiviral therapies may not be able to suppress the progression of tauopathy following SSPE (9). Unlike other conditions, where abundant filamentous tau inclusions are present in all cases (10), only 20% of cases of SSPE have abundant tau inclusions (3,4). It remains to be seen if tau inclusions influence the clinical picture of SSPE. Inflammatory changes also occur in CTE, where microglial cell activation is believed to increase tau pathology and the presence of abundant CD68-positive microglial cells has been demonstrated (38). Moreover, translocator protein (TSPO) positron emission tomography ligands for activated microglia have shown increased signal in retired American football players who are at risk for CTE (39).

It remains unclear how inflammation and microglial cell activation affect tau assembly. Microglial cell activation has been reported to promote tau assembly in mice (40–42) and it also characterizes neurodegenerative diseases with filamentous tau pathology other than SSPE and CTE, the most studied of which is AD (43,44). More work is required to identify the links between neuroinflammation and tau assembly.

## Acknowledgements

This work was supported by the Electron Microscopy Facility of the MRC Laboratory of Molecular Biology. We thank Jake Grimmett, Toby Darling and Ivan Clayson for help with high-performance computing. We also thank Ms R. Otani for technical assistance and Professor B. Ghetti for helpful discussions. For the purpose of open access, the MRC Laboratory of Molecular Biology has applied a CC BY public copyright licence to any Author Accepted Manuscript version arising.

## Author contributions

MH, MT, MS, AA, YI, HM and MY identified the patients and performed neuropathology; MH extracted filaments and performed immunoelectron microscopy and Western blotting; CQ performed cryo-EM data acquisition and structure determination; SHWS and MG supervised the project and all authors contributed to the writing of the manuscript.

## Data availability

Cryo-EM maps have been deposited in the Electron Microscopy Data Bank (EMDB) with the accession numbers EMD-16532, EMD-16535. Corresponding refined atomic models have been deposited in the Protein Data Bank (PDB) under accession numbers 8CAQ, 8CAX. Please address requests for materials to the corresponding authors.

## SUPPLEMENTARY INFORMATION

**Table S1.** Cryo-EM data collection, refinement and validation statistics.

|  | SSPE_case1 |  | SSPE_case2 |  |
| --- | --- | --- | --- | --- |
| **Data collection** |  |  |  |  |
| Microscope | Titan Krios |  | Titan Krios |  |
| Voltage (kV) | 300 |  | 300 |  |
| Detector | Falcon4 |  | Falcon4 |  |
| Magnification | 96,000 |  | 96,000 |  |
| Electron exposure (e-/Å<sup>2</sup>) | 40 |  | 40 |  |
| Defocus range (μm) | -1.0 to -2.0 |  | -1.0 to -2.0 |  |
| Pixel size (Å) | 0.824 |  | 0.824 |  |
| **Data processing** | CTE type I | CTE type II | CTE type I | CTE type II |
| Box size (pixel) | 400 | 400 | 300 | 300 |
| Symmetry imposed | C1 | C1 | C1 | C1 |
| Initial particle images (no.) | 342,441 |  | 388,755 |  |
| Final particle images (no.) | 222,257 | 17,756 | 348,830 | 32,882 |
| Map resolution (Å) | 2.3 | 5.1 | 3.0 | 3.7 |
| FSC threshold 0.143 |  |  |  |  |
| Helical rise (Å) | 2.38 | 2.39 | 2.37 | 2.37 |
| Helical twist (°) | 179.35 | 179.35 | 179.41 | 179.39 |
| **Refinement** |  |  |  |  |
| Model resolution (Å) | 2.9 |  |  | 3.8 |
| FSC threshold 0.5 |  |  |  |  |
| Map sharpening B factor (Å<sup>2</sup>) | -44 |  |  | -111 |
| Model composition |  |  |  |  |
| Non-hydrogen atoms | 2870 |  |  | 3444 |
| Protein residues | 375 |  |  | 450 |
| Ligands | 0 |  |  | 0 |
| B factors (Å<sup>2</sup>) |  |  |  |  |
| Protein | 136.7 |  |  | 259.5 |
| R.m.s. deviations |  |  |  |  |
| Bond lengths (Å) | 0.007 |  |  | 0.009 |
| Bond angles (°) | 1.43 |  |  | 1.746 |
| Validation |  |  |  |  |
| MolProbity score | 1.38 |  |  | 2.49 |
| Clashscore | 1.37 |  |  | 8.40 |
| Poor rotamers (%) | 1.52 |  |  | 4.55 |
| Ramachandran plot |  |  |  |  |
| Favored (%) | 94.52 |  |  | 90.41 |
| Allowed (%) | 5.48 |  |  | 4.55 |
| Disallowed (%) | 0 |  |  | 0 |
| EMDB | EMD-16532 |  |  | EMD-16535 |
| PDB | 8CAQ |  |  | 8CAX |

**Figure S1.**
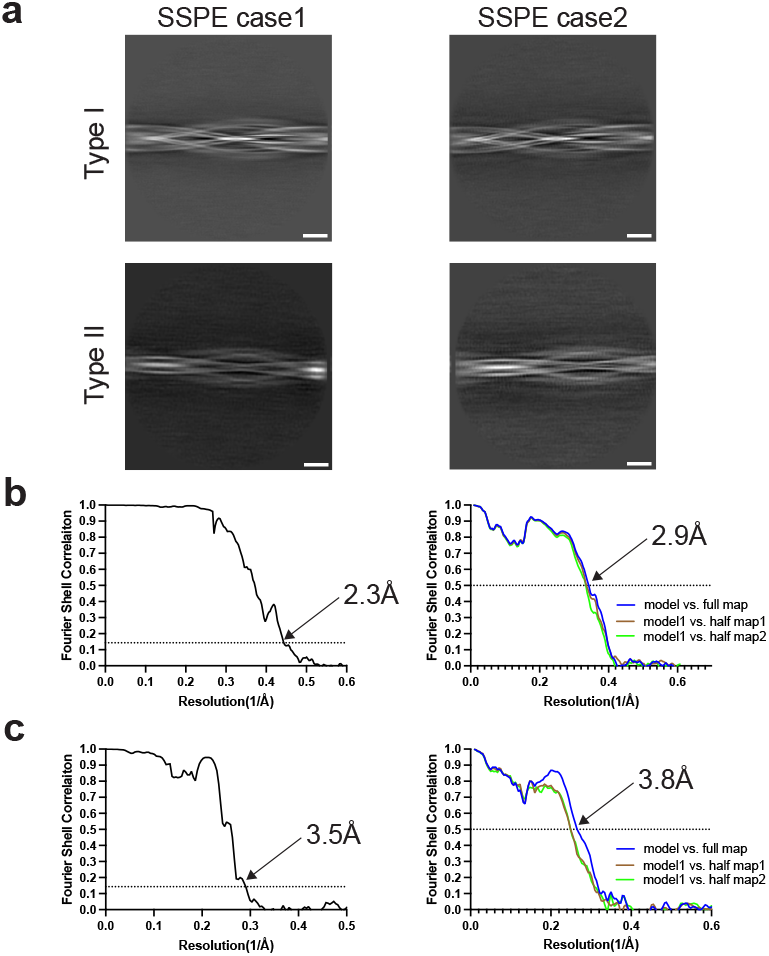
Cryo-EM 2D classifications and Fourier shell correlation (FSC) curves. (a), Representative 2D classification images of tau filaments from both cases of SSPE. Type I and Type II filaments of CTE are in evidence. Scale bar, 10 nm. (b,c), Solvent-corrected FSC curves of cryo-EM half maps (left panel) and model to map validation (right panel). Type I filaments from SSPE case 1 are shown in (b), Type II filaments from case 2 are shown in (c). FSC curves between a model refined in half map 1 and half map 1 are shown in brown (model 1 vs half map 1); FSC curves between the same model and half map 2 are shown in green (model 1 vs half map 2).

